# Spatial and temporal scale-dependent feedbacks govern dynamics of biocrusts in drylands

**DOI:** 10.1101/2025.06.10.658851

**Authors:** Jingyao Sun, Kailiang Yu, Max Rietkerk, Ning Chen, Hongxia Zhang, Guang Song, Xinrong Li

## Abstract

Biota could be ecosystem engineers in generating an intrinsic heterogeneous landscape through scale-dependent feedbacks. Thereby, they can form resource enriched patchiness or islands of fertility, comprising of self-organizing spatial patterns. Research so far has largely focused on the self-organized spatial patterns of plant communities in drylands. It, however, remains unclear whether and how biocrusts having distinct morphology and life history from plant communities could self-organize themselves and form unique spatial patterns. Here we conducted field observations of biocrusts across successional stages and employed a probabilistic Cellular Automaton (CA) model to investigate the distinct self-organized spatial patterns exhibited by mosaic patches of mosses and lichens with different patch size distributions (PSDs). Our study demonstrates that short-range positive feedbacks initially promote the development of patches, featured with a heavy-tailed PSD, while long-range negative feedbacks subsequently curtail further expansion of big patches, thereby establishing a characteristic patch scale with regular PSDs. Strikingly, only lichens reverted back to the heavy-tailed PSD in the late succession stage, presumably implying self-organized critical fragmentation of lichen patches. Field measurements of biocrust performance at the center and edge of patches of varying sizes along succession stages further support the classic scale-dependent feedback mechanism for Turing pattern formation. Collectively, our results clearly demonstrate the capability of the biocrust communities to self-organize themselves to form distinct spatial patterns governed by the spatial and temporal scale-dependent feedbacks, potentially impacting dryland ecosystem functions and resilience.

**Significance Statement:** An intriguing phenomenon is the capability of biota to form striking self-organized spatial patterns. Biocrusts serve as the soil’s skin and have different morphology and life history as compared to plant communities. We investigated these distinct self-organized spatial patterns exhibited by mosaic patches of mosses and lichens across succession stages through field surveys and a probabilistic Cellular Automaton (CA) model. Our results provide empirical evidence that biocrusts act as ecosystem engineers, forming self-organized spatial patterns. Simulations and field measurements of biocrust performance elucidate the mechanism of spatial and temporal scale-dependent feedbacks in governing biocrust spatial pattern formation. This self-organizing ability of biocrusts holds significant implications for ecosystem functions and the resilience of dryland ecosystems.

## Introduction

Patchiness represents a widespread self-organized form of life. It is driven primarily by biota performing as ecosystem engineers in generating an intrinsic heterogeneous landscape (Levin & Paine 1974, Rietkerk et al. 2004, Sheffer et al. 2013). Resource enriched patches or islands of fertility can be formed, comprising of self-organized spatial patterns (Levin 2005, Kéfi et al. 2007a, Franklin et al. 2020). Such self-organized spatial patterns have been widely studied in drylands covering large land surface areas (>40%) and supporting an increasing number of human populations (Schimel 2010, Tarnita et al. 2017, Maestre et al. 2021). Drylands are subject to scarcity of resources (especially water) preventing full homogeneous soil cover by biota. The scale-dependent feedback interacting with biota (i.e., vegetation) that perform as ecosystem engineers facilitates growth in short ranges through positive feedback (i.e., regulated via vegetation and rainfall infiltration) while competing through negative feedback in long ranges, thus constituting scale-dependent feedback (Klausmeier 1999, Tarnita et al. 2017, Rietkerk et al. 2021). This scale-dependent feedback mechanism leads to the formation of diverse Turing patterns of plants, such as spots, labyrinths, gaps, or fairy circles (Pringle and Tarnita 2017, Franklin et al. 2020). Research in the previous decades has largely focused on theoretical aspects of the self-organized spatial patterns formation in plant communities (Klausmeier 1999, Gilad et al. 2004, Rietkerk et al. 2021). However, empirical evidence of the underlying mechanisms of scale-dependent feedback and pattern formation remained limited (but see Scanlon et al. 2007, Kéfi et al. 2007a, Xu et al. 2015). Recent studies have shown that drylands dominated by biocrusts represent one of multiple possible stable states and have underscored the role of biocrusts as landscape-shaping ecological engineers (Chen et al. 2020). Therefore, there is a pressing need to bridge the knowledge gap between theoretical foundations and empirical validation of self-organization processes within biocrust communities.

Biocrusts, comprising non-vascular plants and photoautotrophs such as lichens, mosses, cyanobacteria, along with various heterotrophic microbes, constitute the primary soil surface cover and act as soil skin in arid lands (Li et al. 2016, Weber et al. 2016). It largely regulates the biogeochemical cycling and serves as interface between atmosphere and soil (Porada et al. 2013, Weber et al. 2022). Biocrust patches can establish local feedbacks through their interactions with the surrounding environment, such as through water usage or regulation of the water balance, potentially leading to the formation of self-organized spatial patterns. (Turnbull et al. 2012, Kinast et al. 2013, Eldridge et al. 2020). These biocrusts typically develop on bare ground through successional stages (i.e., from the early dominance by cyanobacteria) to finally form a climax community dominated by mosses or lichens. Previous research on the spatial development of biocrusts has primarily focused on cover at equilibrium, often assuming that spatial distribution is a direct reflection of environmental changes, with limited empirical evidence on the ongoing process of self-organization or transient states (Heras et al. 2011, Kéfi et al., 2007a, 2007b, Xu et al., 2015).

During the self-organization process of biocrust patches, the dominance of positive and negative feedback mechanisms shifts across different temporal stages, indicating an asynchronous nature in scale-dependent feedbacks (Hastings 2016). Indeed, at the initial expansion of biocrust patches on barren soil, the large patches are endowed with a growth advantage and such expansion (i.e., sustained by short-range positive feedback) would result in a heavy tail in the PSDs. However, subsequently a shift to long-range negative feedback due to competition for limited space or resources could restrict further expansion of large biocrust patches, thus ultimately leading to a characteristic scale in patch size and a regularity in spatial pattern (von Hardenberg et al. 2010). The relatively rapid dynamics and restricted spatial scales of biocrust patches may offer a unique opportunity to study the asynchronous and scale-dependent dimension of the self-organization process in sessile life forms, which is often challenging to observe in plant communities due to its slow and longtime life history (Bowker et al. 2014, Nelson et al. 2022).

Biocrusts of lichens and mosses are characterized by different morphology, life history, and ecological roles, and form complex and distinctive patch-interpatch mosaics shaping ecosystem functions (Berdugo et al. 2017, Bowker et al. 2013, Bowker et al. 2010, Sun et al. 2021b). Positive feedback is often observed in established patches, particularly the larger ones. This can be attributed to factors such as improved water absorption and soil moisture retention from the aggregation of moss stems, or enhanced soil stability for lichens due to the aggregation of thalli (Belnap and Büdel 2016, Sun et al. 2021a). This local positive feedback mechanism drives the expansion of both lichens and mosses, albeit through these different facilitation pathways. Moreover, contrasting life histories—lichens’ high tolerance of low-resource environments versus mosses’ rapid growth in wet conditions—lead to spatially distinct self-organization trajectories (Read et al. 2016). For instance, mosses with their developed hydraulic conductivity system often outcompete lichens in the later stages of succession when conditions become more favorable (Sun et al. 2021b). Previous studies have observed the successional replacement of lichens by mosses, inevitably influencing their self-organizing trajectories (Sun et al. 2021b, Weber et al. 2016). As such, the contrasting yet simple morphologies of biocrust components—lichen versus moss—enable the display of scale-dependent characteristics on the soil surface without vertical stratification of their physical structures. Indeed, while pattern formation has been extensively studied in vegetation with focus on either woody plants or grasses (Klausmeier 1999, Rietkerk and Van de Koppel 2008, Rietkerk et al. 2021), the relatively simple structure of biocrusts may offer greater convenience for manipulation and observation of mosaic patterns by accounting for the idiosyncratic features of lichens and mosses (Bowker et al. 2014), thereby potentially revealing fundamental mechanisms that are universally applicable.

Here we adopted an integrative methodology comprising field observations and computational modeling to examine the self-organized spatial patterns of mixed biocrust communities – lichens and mosses. Based on the stock-flow framework, an increase in patch size (stock), that enhances resource flow to the patch, accelerates further accumulation of patch, creating positive feedback. Conversely, an increase in stock that inhibits flow stabilizes the stock, forming negative feedback. Thus, we formulated the hypotheses (Fig. 1A) that local propagation predominantly facilitates the growth of large biocrust patches through short-range positive feedback during the initial colonization of vacant habitats, consequently resulting in a heavy-tailed PSD. As patch density increases, resource constraints hinder further patch growth due to long-range negative feedback in over-crowded environments, leading to a limited number of big patches and the emergence of a characteristic scale in PSDs. To test these hypotheses, we used field observations derived from a chronosequence of biocrust succession during six moments in time (each surveyed with 60 one-dimensional transects of 1.5 m to investigate the temporal change in PSD between lichen and moss. Moreover, we introduced a probabilistic Cellular Automaton (CA) model (Fig. 1B), incorporating simple update rules of patch propagation, mortality due to resource constraints, dispersal, and spatial competition between lichens and mosses, to represent the governing positive and negative feedback mechanisms of patch growth for a mechanistic understanding of the self-organization of biocrusts. Further, we analyzed the performance of lichens vs mosses at various developmental stages to elucidate the local mechanisms underlying the observed global pattern.

**Figure 1.**
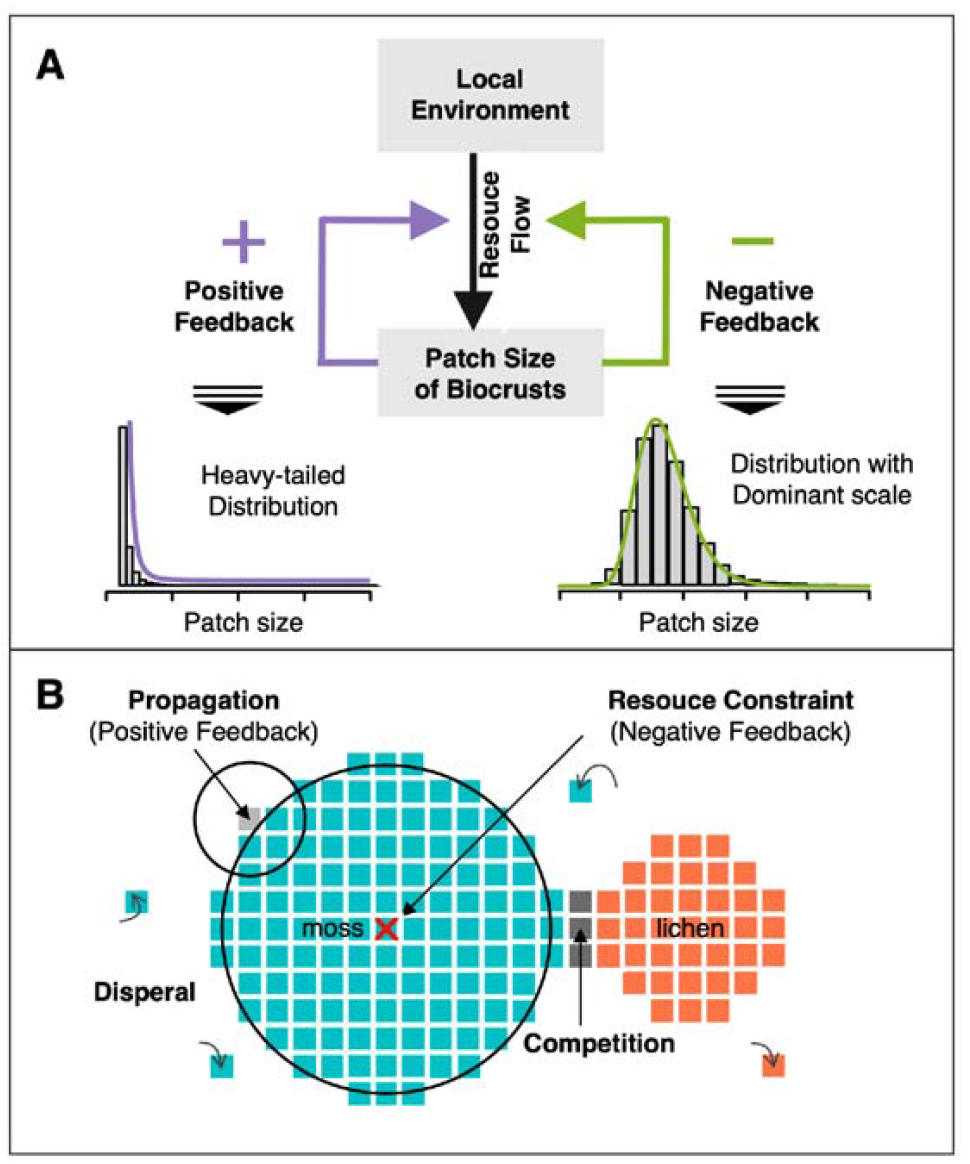
Conceptual and computational framework of self-organized biocrust spatial pattern. A) Conceptual schema of the working hypothesis. Grey boxes and broad arrows depict the stock-flow relationship: an increase in stock (patch size) that promotes flow (resource) accelerates further accumulation, creating positive feedback and heavy-tailed PSDs. Conversely, an increase in stock (patch size) that inhibits flow (resource) stabilizes the stock, forming negative feedback and PSDs with a characteristic scale. In PSDs, the x-axis represents patch size, while the y-axis typically represents frequency or density. B) Cellular automaton model rules governing biocrust patch dynamics. **Propagation:** Vacant cells adjacent to patches (grey cells) have the potential to be colonized. Cells in adjacent range (small circles) have enhanced colonization rates, constituting short-range positive feedback. **Resource constraint (death)**: A cell’s death (red cross) is triggered when the local density (large circle around the cross) surpasses the carrying capacity, inducing long-range negative feedback. **Dispersal**: Vacant cells may sporadically become colonized by biocrusts with a certain probability. **Competition**: When both moss and lichen potentially encounter in a cell, mosses are more likely to win. The propagation rule prioritizes the growth of large patches (positive feedback), leading to heavy-tailed PSDs. Resource constraints impede the further growth of patches in overcrowded local environment (negative feedback), resulting in PSDs with a characteristic scale.

## Results and discussion

### The emergence of different patch size distributions

Our field observations indicate a succession in biocrusts accompanied by significant changes in PSDs. Cyanobacteria and algae achieve their highest coverage at 13 years; lichens’s coverage peaks between 23 and 36 years; and mosses increase in coverage over time (Fig. 2A). We employed power-law relations (PLR) to quantify the heavy-tailed characteristics of the patch size distribution. This distribution is manifested as an irregular pattern, marked by numerous small patches interspersed with a few large ones. In contrast, regular patterns exhibit a more uniform configuration, with most patches displaying similar sizes. Figure 2B illustrates this contrast. In the early stages of succession, patch size and frequency follow a PLR with moss or lichen patches exhibiting the scaling exponents α greater than 2 (Fig. 2E, F). When converting one-dimensional patch scales to two-dimensional scales, the scaling exponent is divided by a factor of 2. We arbitrarily set it at 1.5, considering the fractal shape of patches. Interestingly, the scaling exponent of PLR in biocrust communities, which falls within a proxy range of 2 to 3 indicative of scale-free characteristics (von Hardenberg et al. 2010), is generally consistent with previous findings on dryland vegetation (Sankaran et al. 2019, Scanlon et al. 2007). Furthermore, the smaller scaling exponent in moss PSD compared to lichens suggests stronger positive interactions among mosses in generating relatively bigger patches, possibly due to a vertical structure that facilitates direct nursing effects (Sun et al. 2021a, b). In contrast, the positive feedback in lichens (primarily from thalli aggregation) is weaker, thus resulting in a relatively smaller number of large patches with larger scaling exponent values than moss (Belnap and Büdel 2016).

**Figure 2.**
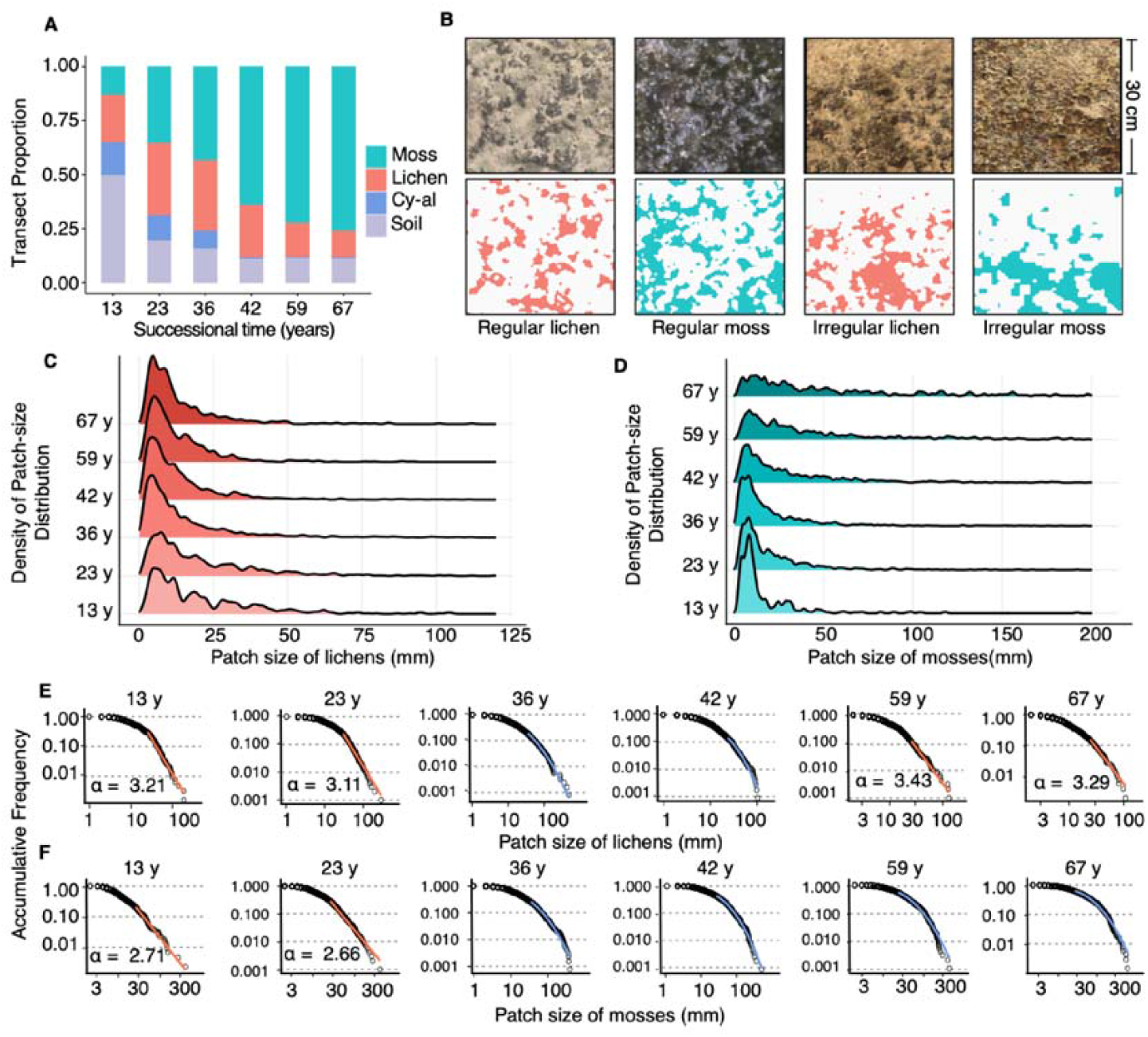
Dynamics of patch size distribution of biocrust communities along gradient of succession (spatial development). A) Changes of transect proportion, indicating shifts in biocrust coverage and composition over time during succession. B) Field observations reveal irregular patterns that indicate heavy-tailed PSDs, characterized by the blend of numerous small patches and a small number of disproportionately large patches. Regular patterns suggest a more uniform distribution where most patches are of comparable size, indicating a characteristic scale within the PSDs. C-F) Patch size distributions (PSDs) from field observations across the chronosequences. Upper subfigures (C, D) represent density of patch size distributions for lichens (red) and mosses (green), respectively. Lower subfigures (E, F) show reversed cumulative frequency of lichens and mosses patches on a log-log scale. Power-law fits are marked in red if it is significant according to the Kolmogorov-Smirnov (KS) test. Otherwise, blue log-normal curves depict the general trend without PLR.

During the subsequent stages of moss and lichen PSD dynamics, small and large patches tend to develop towards a medium size and ultimately a deviation from the power-law distribution is evident for both functional groups. The log-log accumulative frequency of PSDs assumes a convex shape, with inflection points around 50 mm for lichens and 100 mm for mosses, indicating the emergence of a characteristic patch scale. Model results indicate that the final stable patch size is related to the range of negative feedback. In this study, we did not explicitly set how biocrusts utilize resources but implicitly reflected their utilization of surrounding resources by constructing local density. Mosses, compared to lichens, with their vertical structure and initial differentiation of stems and leaves, can generate a larger hydraulic gradient and exploit resources over a larger range, resulting in a larger range of negative feedback and, consequently, larger patches (Bowker et al. 2014). This distribution also highlights the increasing regularity during succession (Fig. 2B). These results are generally consistent with the principle of “Turing patterns”, whereby a heavy tail in PSD forms initially with the dominance of short-range positive feedback, which is gradually displaced with the dominance of long-range negative feedback (Pringle and Tarnita 2017, Rietkerk and van de Koppel 2008). Collectively, our results suggest that biocrust patch propagation and long-range resource constraints are sufficiently universal for emergence of Turing patterns through scale-dependent feedbacks.

Notably, the distribution of lichen patches reverts to a power-law in the late succession stage (Fig. 2 E). Indeed, unlike the free growth observed in the early stages of succession, patch growth at late stage is constrained due to space limitations. This constraint limits the number of big patches by mosses, thereby leading to deviation of PSD from PLR, also increased the environmental stresses (i.e., by competition from moss) for lichens. Therefore, the intriguing phenomenon of lichens reverting to PLR can likely be attributed to competition and degradation. As lichens begin to decline, external pressures place them in critical state whereby positive feedback maintains the superiority for large patches. Fragmentation initiates for smaller patches and these are more likely to continue breaking down until they can avoid interference from mosses. Consequently, mosses gain space, enabling them to adjust their spatial configuration and potentially causing further fragmentation among remaining lichen patches. Collectively, these processes result in the formation of numerous small lichen patches alongside a few unaffected large ones, re-creating a heavy-tailed distribution. Previous studies have frequently highlighted the connection between self-organized criticality and PLR (Kefi et al. 2007, Verwijmeren et al. 2013). The cascading adjustment of lichen patches described above aligns with the changes that occur during self-organized criticality, where an initial change can trigger a chain reaction, resulting in a dynamic shift in the ecosystem’s structure (Muñoz 2017). Therefore, it is suggested that the return of lichen PSD to PLR may represent a self-organized critical phenomenon. During the stage of overall lichen patch fragmentation and degradation, lichen patches may maintain system resilience and prevent system collapse through self-organization.

### Changing feedbacks account for the spatial pattern dynamics

The results of the simulations closely align with field observations, validating the working hypothesis of asynchronous expressions of scale-dependent feedback and phased development in the spatial organization of biocrust communities (Fig.1). This developmental process is initially characterized by the “patch growth phase” (Fig. 3A, S3). During this phase, notable rapid increases in density of both mosses and lichens are observed alongside PSDs that not only exhibits a pronounced heavy tail in PSD (as seen in Fig. 3 B, F) but also undergoes progressive tail elongation with subsequent iterations (Fig. 3 C, G). The positive feedback mechanism in the early stage parallels the classic “preferential attachment” principle, which describes a “rich-get-richer” phenomenon in non-equilibrium dynamics (Barabasi et al. 1999). As densities of mosses and lichens approach the environmental carrying capacity, an optimization in the spatial arrangement of patches occurs-the ‘pattern formation phase’. This optimization manifests as a transition from irregular to regular spatial configurations of both lichens and mosses (Fig. 3 A, snapshots 1-2). This phase involves the fragmentation of exceptionally large patches and the merging or continued expansion of smaller patches, all converging towards a characteristic scale, as evidenced in Fig. S3. This pattern aligns with our proposed hypothesis, highlighting the decisive role of negative feedback in determining the size and scale in patches (Fig. 1). Despite the asynchronous expression of positive and negative feedbacks in determining patch size; however, competition between mosses and lichens also influences the subsequent PSDs, as shown in the patch fragmentation phase (see the grey area of Fig. 3A). During this phase, lichen patches fragment rapidly and then re-form irregular patterns without a characteristic scale, indicating that the replacement of lichens by mosses represents a critical transition.

**Figure 3.**
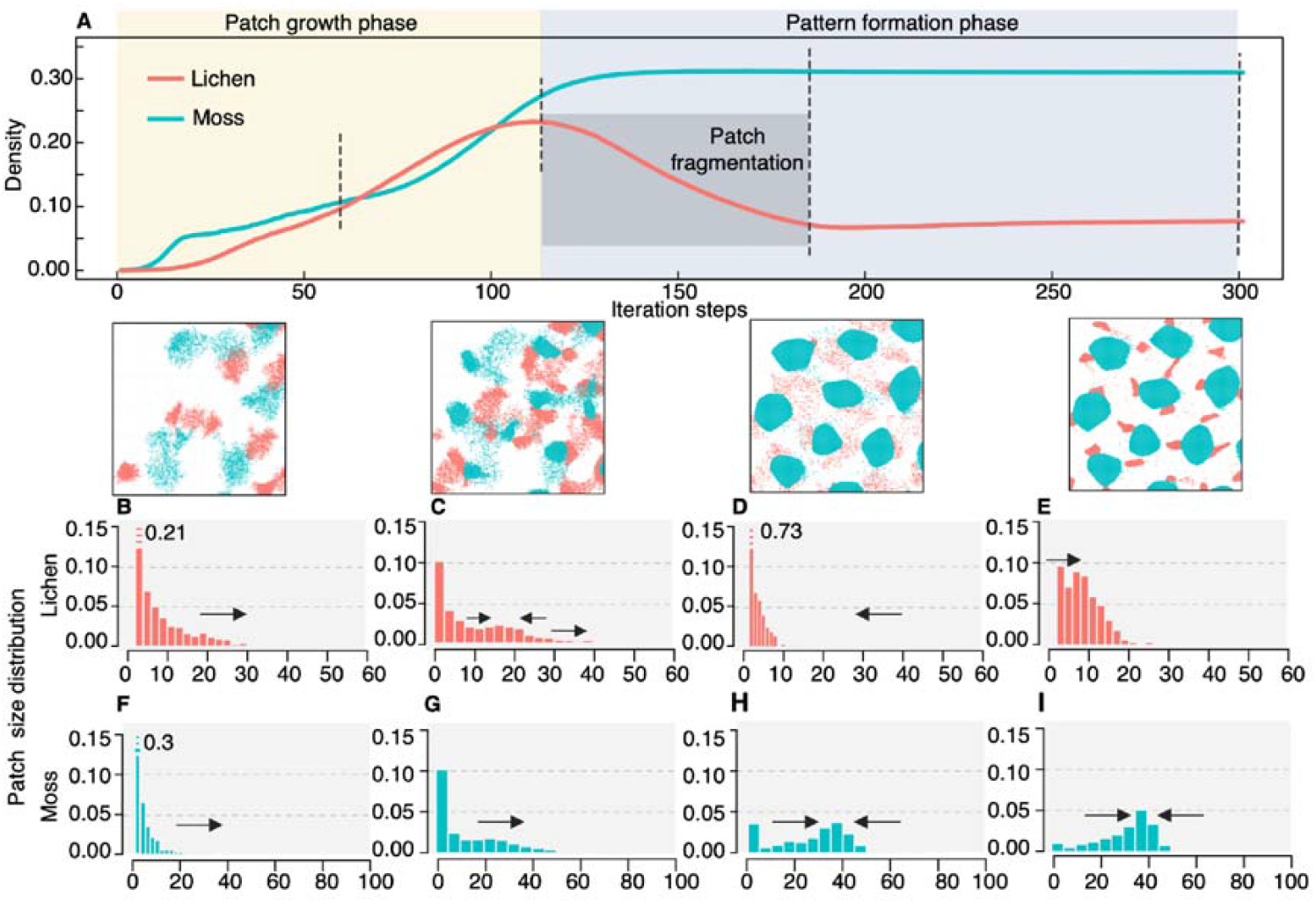
Simulated dynamics of lichen and moss patches across succession. A) Depicts the variation in cellular density with iteration steps. Solid lines illustrate the density changes under these trajectories. The snapshots below the graph show the spatial distribution at specific iterations (marked by black dashed lines). The pale-yellow area delineates the patch growth phase, focusing on the primary occupation process of patches in vacant habitats. The light purple area delineates the pattern formation phase, characterized by minimal density changes and the emergence of regular patterns. The grey area indicates the patch fragmentation phase in lichens, during which the characteristic scale in the PSD vanishes. B-E) and F-I) present the frequency distribution histograms of patch distributions at the marked iteration steps for lichens and mosses, respectively, elucidating the spatial and quantitative dynamics of patch formation and evolution across the iterations. Arrows denote the direction of change in frequency distribution across iteration steps.

In summary, our theoretical modeling results underscore the decisive impact of self-organization based on local feedback on the compositional changes in biocrust community. Traditional studies, by contrast, often implicitly assumed each stage of biocrust development to be in equilibrium with gradually changing environmental conditions (Housman et al. 2006) and made causal inferences based on this premise. However, our findings suggest that, given specific environmental conditions, the biocrusts go through several successional stages in time rather than that they remain in a stable equilibrium. This indicates the need for future studies to employ more non-equilibrium research methods, especially when time is involved (Klaus and Rohde 2005).

### Biocrust performance reflect emergent scale effect

The results of biocrust performance in the center vs edges of the patches across succession stages further support the effect of an emergent scale. In the early succession stage, independent of patch size, growth within the centers of biocrust patches generally outperforms that at the edges. As such, the performance of biocrusts in larger patches is generally superior to that in smaller patches in the early stage of succession (Huang et al. 2023). This is evidenced by higher chlorophyll a and b content in lichens and higher biomass, density, and height in mosses (Fig. 4B-F). Chlorophyll measures lichen biomass, revealing significantly greater growth within patches than at the edge (Fig. 4 B, C). Moss biomass adheres to the same trend, with individual density and height indicating that this elevated biomass is a result of contributions from both large individuals and dense population (Fig. 4 D-F). This biomass enhancement underscores the system’s propensity for rapid expansion driven by positive feedback mechanisms. This better performance in the center during early succession may be attributed to either the nursing effects from established neighbors or an inward-to-outward growth form (Sun et al. 2021b).

**Figure 4.**
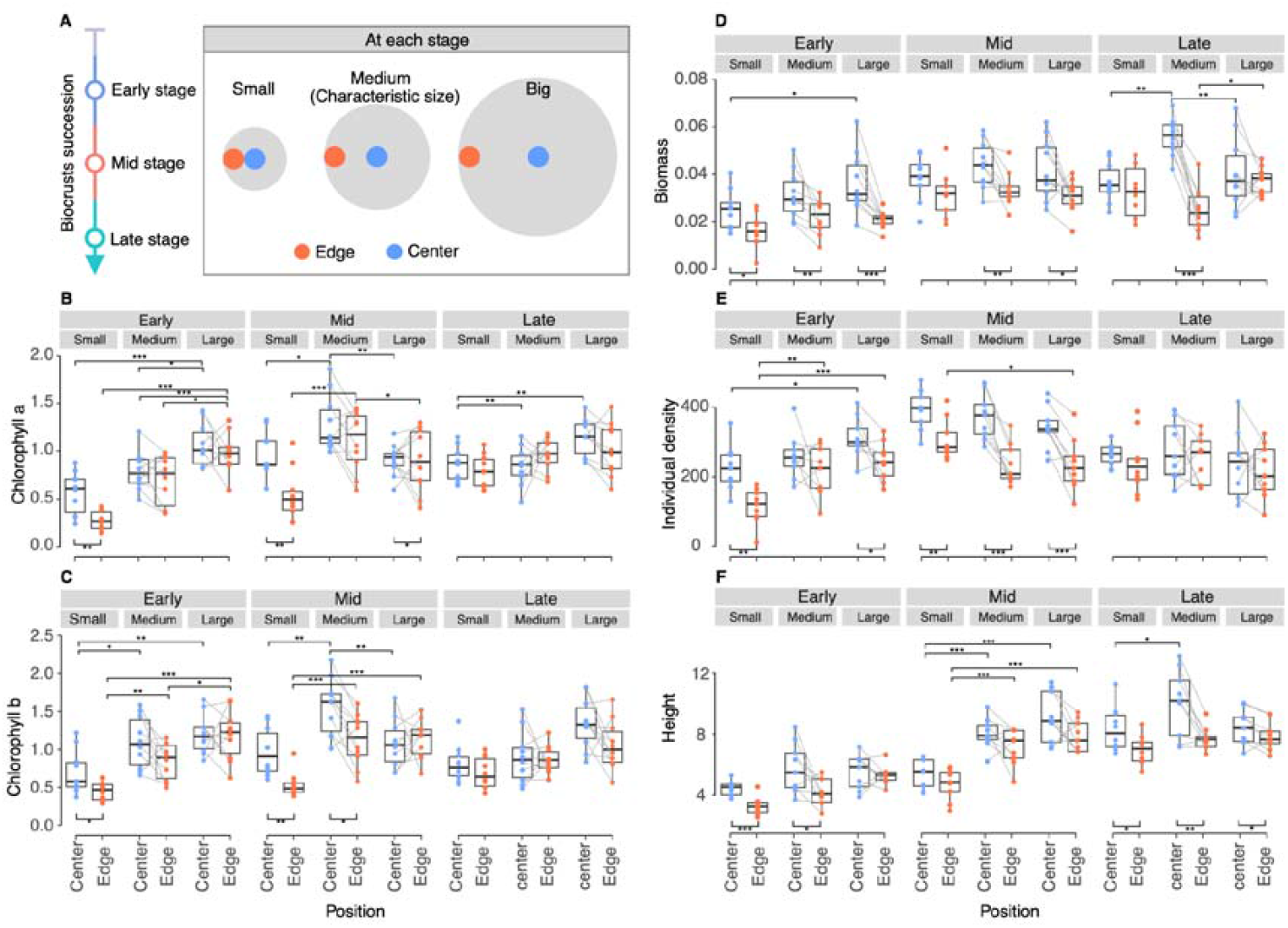
Growth variability in lichen and moss between biocrust patch centers and edges across succession stages. A) Schematic sampling for growth performance of moss and lichen. The central-edge differences are compared across various patch sizes (small, medium, large) and at different stages of ecological succession (early, mid, late). B, C) Variations in chlorophyll a and b content in lichens. D, E, F) Variations in moss biomass, stem density and height. Significance markers below the box plots compare central-edge differences; for small patches, where simultaneous center and edge sampling was not possible, a t-test was used. For medium and large patches, gray lines connect center and edge measurements within a patch, with paired t-tests applied. Significance markers above the box plots compare growth performance across different patch sizes at the same patch location, using pairwise multiple comparisons based on one-way ANOVA. Significance levels: * p < 0.05, ** p < 0.01, *** p < 0.001.

Negative feedback also influences patch-scale biocrust performance (Fig. 4B-F), alongside the emergence of characteristic scales in PSDs (Fig. 2 E F). For lichens, the PSDs reveals a characteristic scale during the mid-successional stage (Fig. 2 E), which corresponds to the significantly higher chlorophyll content in intermediate-sized patches compared to small and large ones (mid panel in Fig. 4 B, C). In contrast, the early stage shows a significant higher chlorophyll content in large patches (left panel in Fig. 4B, C). These findings suggest the substantial inhibitory effects of long-range negative feedback. Furthermore, the advantage of medium-sized lichen patches dissipates in the late successional stage, with no significant central-peripheral differences (right panel in Fig. 4B, C). This supports the explanation of critical transition of lichen, associated with patch fragmentation, underlies the PLR observed in lichen PSDs during the late successional stage (Fig. 2 E). Concurrently, fragmentation blurs the distinction between patch centers and edges. For mosses, medium-sized patches gain an advantage in the late successional stage, with center biomass significantly higher than in small and large patches (right panel in Fig. 4D) and height significantly greater than in small patches (right panel in Fig. 4F). This superior performance of medium-sized patches also coincides with the emergence of characteristic scales in PSDs (Fig. 2F). Results also show a decline in individual density (Fig. 4 E) alongside an increase in individual height (Fig. 4 F). These results, suggest that biomass accumulation stems from enhanced individual performance (Fig. 4 F) rather than population size (Fig. 4 E), which may have decreased due to self-thinning. This self-thinning also indicates the role of negative feedback in patch dynamics at this stage.

While the theoretical aspects of self-organized spatial patterns have been widely clarified in terms of scale-dependent feedback in dryland vegetation communities, the empirical observations on the underlying mechanisms remain scarce (i.e., with limited observations of resource patches or islands of fertility). Here our study uses a patch-scale evaluation of biocrust performance through empirical evidence to show the varying growth performance in center vs edge across small, medium, and big patches along the succession stage in dryland biocrust communities. Our results collectively demonstrate the patch expansion trend from center to edge and the superiority of the medium-sized patch with the establishment of regular patterns, thus supporting the potential impact of spatial patterns on ecosystem function. Given the significant role of trait-based performance, which can be easily measured in the field, we suggest increasing field measurements of localized performance and/or incorporating them into theoretical models to enhance our understanding of self-organized biological spatial patterns.

### Implication of self-organized biocrust spatial pattern

Our study uses an integrative approach by combining field observations and a probabilistic CA model to clearly demonstrate the development of self-organized spatial patterns in dryland biocrust communities. It offers novel insights with significant implications for dryland life organization and ecosystem functions and resilience. First, our results demonstrate the capacity of biocrusts as ecosystem engineers to spontaneously self-organize and form Turing patterns. This ability can lead to environmental heterogeneity, thereby influencing the functions and resilience of dryland ecosystems (Bowker et al. 2013, Kéfi et al. 2024). Biocrusts, as ecosystem engineers, have unique characteristics due to their small sizes of individuals and fast responses as compared to plants, thus suggesting their unique role in self-organizing themselves and potentially and uniquely influencing the ecosystem functions and resilience across dryland landscapes. Second, our study demonstrates the temporal change of self-organized biocrust spatial pattern in which PLR initially form but gradually disappear during succession stage. This result highlights the temporal dimension of self-organized spatial patterns (Hastings 2016), while previous studies have largely used the spatial data on stable or climax communities to examine the pattern formation process (Kéfi et al. 2007a, Sankaran et al. 2019, Scanlon et al. 2007). Third, our study displays the intriguing self-organized spatial patterns in the moss-lichen mosaic pattern. This pattern likely infers the unique and distinct moss vs lichen morphology and life history, thus highlighting the importance of accounting for moss vs lichen interaction/competition for a comprehensive understanding of biocrust spatial patterns. Fourth, our field measures of biocrust performance uncover the underlying mechanisms in terms of the growth pattern in center vs edge of biocrust patches across patch size along the succession stage to collectively support the classic scale dependent feedback in pattern formation. This performance or trait perspective, which has been previously shown to largely influence community assembly and functions (Concostrina-Zubiri et al. 2018), offers new opportunities to scale up from traits, and individual performance to patch expansion or self-organized spatial patterns in future studies.

### Conclusion

Here we have proposed a minimal conceptual framework grounded in the interplay of positive and negative feedback and combined empirical observations with a theoretical model to elucidate the self-organized spatial patterns in dryland biocrust communities. The positive feedback leads to an irregular biocrust patch distribution with heavy-tailed PSD when global density is low, aligning with the principles of preferential attachment. As patches grow, negative feedback induced by resource constraints shapes pattern formation, achieving global optimization of resource utilization. The emergence of this regular pattern with a characteristic patch size not only shape the spatial configuration of biocrust patches but also influences the living performance of biocrusts, thus, potentially influencing ecosystem functions. This presents a valuable perspective for exploring the connections between self-organization and ecosystem functionality. The field measures of biocrust performance collectively support the classic scale-depended feedback mechanism in invoking spatial pattern formation, which has the implications of leveraging localized performance for a better understanding of pattern formation.

## Materials and Methods

### Field observations

Field observations were conducted in Shapotou, located within the Ningxia Hui Autonomous Region, on the southeastern fringe of the Tengger Desert (37°32′-37°26′ N, 105°02′-104°30′ E, 1300-1350 m above mean sea level). This site is characterized as a typical ecotone, transitioning from desertified steppe to sandy desert (Li et al. 2003). Meteorological data from the local station (SDRES, CAS) spanning 1955 to 2016 indicate an annual mean temperature of 10.0 °C and an annual mean precipitation of 186 mm. The prevalent soil type in this area is wind-borne sand, classified as *Eutric Arenosols* in the World Reference Base. In this study, the research was conducted on a series of sand-binding vegetation belts which were established directly on sand dunes at various intervals, specifically in the years 1956, 1964, 1981, 1987, 2000, and 2010. This systematic establishment has resulted in a revegetation chronosequence, providing a temporal record of the spatial progression of biocrust development. Further details on the study area can be found in the supplementary material (Fig. S1).

We employed the space-for-time substitution method to analyze ecological succession. The survey was conducted in 2023, with different sand-binding vegetation belts corresponding to 13, 23, 36, 42, 59, and 67 years of biocrust spatial development. To evaluate the community composition of biological soil crusts (biocrusts), unidimensional interception length measurements conducted along line intercept transects were adopted, hereafter referred to as “transects” (Bowker et al. 2010). For each established sand-binding belt, forty transects, each measuring 1.5 meters in length, were systematically laid out. In consideration of spatial heterogeneity, these transects were positioned at intervals of no less than 2 meters apart. To mitigate potential confounding influences from vegetation, litter, and micro-geomorphological features, the transects were deliberately situated on flat terrain, maintaining a minimum distance of 30 cm from the nearest shrub or tussock and at least 0.5 meters from the nearest soil mound (Choler et al. 2001).

Throughout the length of each transect, we meticulously recorded the presence of various biocrust constituents – including mosses, lichens, cyanobacteria, algae, rocks, plants, and bare soil – at every 1 mm interval, achieving a resolution of 1-mm. This resolution was deemed adequate for our study as the smallest observed BSC patches exceeded 1 mm in size. Mosses and lichens were identified to species level in situ, whereas cyanobacteria and algae were collectively categorized as a single “species” due to the impracticality of field identification. Consequently, each transect yielded a dataset comprising 1500 records, represented as a sequence (e.g., “MMMMLLLMMMM…”). The total count of a particular letter, or “taxon”, was considered indicative of its abundance. Furthermore, each contiguous string of identical letters within the sequence was designated as a “patch”.

### Patch size distribution

We used the power-law (PL) fitting of the transects data to evaluate the heavy-tail properties of PSDs in natural biocrusts.

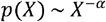

In this probability density function, X represents the transect length of a patch. Only patches with lengths greater than, X_min_ are included in the power-law fitting. The scaling exponent α characterizes the shape and convergence of the distribution. For our two candidate models, we empirically set X_min_ to 20 mm or 30 mm when fitting observed lichen/moss patches, because patches smaller than X_min_ are fragile and do not reflect the dynamics of patch growth. The X_min_ for simulated patches was identified by minimizing the KS distance between the fitted model and data. The exponent of the distribution was estimated using Maximum Likelihood Estimation (MLE), and the parameter uncertainty was determined by bootstrap. The fraction of synthetic datasets that result in a fitted model with a K.S distance larger than the K.S distance calculated when fitting our dataset, was considered the p-value of our fit. A p-value above 0.1 represents a good fit. The model fittings were performed with “poweRlaw” package on R platform (Gillespie 2015).

### Cellular automaton

The probabilistic CA models were used to simulate the growth of biocrust patches. The CA model facilitated simulations of both single-component patch growth, to validate the positive/negative hypothesis (Fig. S3), and dual-component simulations (moss and lichen) to replicate the PSD changes observed in field studies. In CA model, each cell within the model was assigned a specific state (e.g., bare soil, lichen, or moss) and grew in a 2-D null lattice (400 × 400) with vanishing boundary. The main principle of state transition is determined by the following four process: local propagation, dispersal, resource constraint (leading to death), and competition (refer to Fig. 1 B):

### Dispersal

Vacant cells may sporadically become colonized by biocrusts with a low probability, accounting for air-borne debris or biocrust propagules.

### Propagation

Vacant cells adjacent to existing patches possess the potential for colonization. The likelihood of occupancy is positively correlated with the number of neighboring cells, a neighboring facilitation determined by an adjacent range (small circle in Fig. 1 B) surrounding the focal cell. This correlation indicates the intensity of the short-range positive feedback mechanism. **Resource constraint (death):** If a site is occupied by many cells sharing resources, cell death is triggered when local density exceeds the local carrying capacity. Local density is calculated by dividing the number of occupied cells by the total area within the local range (big circle in Fig. 1 B).

### Competition

In scenarios where both moss and lichen potentially encounter in a cell, mosses are more likely to prevail in spatial competition. This model assumes direct competition rather than indirect exploitation, given that lichen and moss have independent carrying capacities.

To reflect changes in the soil environment due to succession (Li et al. 2003), the local carrying capacities for lichens and mosses were modeled following unimodal and saturation trajectories, respectively (Fig. S2, Table S2). For compatibility with field data, patch sizes (cell segment) were recorded from 100 simulations using a line transect method akin to field evaluations. Details and parameterization of the model can be found in appendix S1.

### Performance indicators

This study aims to compare growth differences of lichen and moss between the centers and edges of various biocrust patches, focusing on the impacts of succession time and patch size. Sampling was conducted during early (13 years), mid (36 years), and late (67 years) successional stages. Three levels of patch sizes were selected: Small patches: approximately 2 cm in diameter or smaller. Medium patches: consistent with the characteristic scale observed in the late stage, approximately 4 cm in diameter for lichen patches and about 5 cm for moss patches. Large patches: 10 cm or larger in diameter. Sampling was conducted using a 3 cm diameter, 3 cm high PVC ring, collecting on the upper biocrust layer to the laboratory for further analysis. Patch selection was non-random, performed in well-developed, typical areas, ensuring environmental homogeneity by excluding vegetation, mounds, tussocks, and bare patches. Only near-circular patches were chosen, with 9 replicates for each treatment group.

The lichen performance focused on chlorophyll a and b content. A 6.58 mm diameter (0.34 cm^2^) metal ring sampler was used to collect samples from marked positions in the PVC ring centers. To each soil sample, 10 mL of ethanol (99.9%) was added in a 50-mL cap tube, then heated for 5 min in an 80°C water bath. After cooling for 30 min, the mixture was centrifuged at 4000 rpm for 5 min. Light absorption of the extracted solutions was measured at red absorption peaks of wavelengths of 665/649 nm using a spectrophotometer (UV-1700 PharmaSpec, Japan). The results were then converted to μg/cm^2^ based on the sampling area. For mosses, performance indicators including biomass, stem density, and stem height were assessed. The soil was first washed off, stem numbers in 0.34 cm^2^ metal ring were counted manually, and average height was measured. Samples were then dried in an oven at 65°C for 48 hours and biomass measured. For small patches, where simultaneous center and edge sampling was not possible, the center-edge difference of indicators was evaluated via t-test. For medium and large patches, paired t-tests applied. The differences in performance indicators between patch size were evaluated using analysis of variance (ANOVA).

## Acknowledgments

This work was supported by the National Natural Science Foundation of China (grant nos. 32401449 and 32061123006). The research of MR is supported by the European Research Council (ERC-Synergy project RESILIENCE, proposal nr. 101071417) and by the Dutch Research Council (NWO ‘Resilience in complex systems through adaptive spatial pattern formation’, project nr. OCENW.M20.169).

