## Supplement materials for "Spatial and temporal scale-dependent feedbacks govern dynamics of biocrusts in drylands"

*Kailiang Yu

*Xingrong Li

**This PDF file includes:**

Supporting text

Figures S1 to S3

Tables S1 to S2

**Overview of the study area**

Field observations were conducted in Shapotou, located within the Ningxia Hui Autonomous Region, on the southeastern fringe of the Tengger Desert (37°32′-37°26′ N, 105°02′-104°30′ E, 1300-1350 m above mean sea level). This site is characterized as a typical ecotone, transitioning from desertified steppe to sandy desert. Meteorological data from the local station (SDRES, CAS) spanning 1955 to 2016 indicate an annual mean temperature of 10.0 °C, with the mean temperatures for January and July being -6.9°C and 24.3°C, respectively. The region receives an annual mean precipitation of 186 mm, predominantly between May and September (accounting for 80% of the total). The annual mean wind velocity is 2.9 m/s, and the potential evaporation rate is approximately 2900 mm. The prevalent soil type in this area is wind-borne sand, classified as *Eutric Arenosols* in the World Reference Base.

To protect the Baotou-Lanzhou railway, a series of sand-binding vegetation belts were established directly on sand dunes, utilizing shrubs such as Caragana korshinskii Kom and Artemisia ordosica Krasch, along with straw checkerboards (Fig. S1). These belts, each extending 500 m in width along the railway, were developed in a chronosequence (1956, 1964, 1981, 1987, and 1990). The 1990 belt was excluded from our study due to recent artificial disturbances from tourism. Instead, we focused on a nearby (1.3 km) sand-binding belt constructed in 2000 and 2010, located at the Soil Water Balance Experimental Field of the Shapotou Desert Experimental Research Station, Chinese Academy of Sciences. The survey was conducted in 2023, with different sand-binding vegetation belts corresponding to 13, 23, 36, 42, 59, and 67 years of biocrust succession. These belts, selected for their uniform climatic conditions and identical sand-binding treatments, represent various successional stages of sand-binding vegetation and biocrusts. Post-revegetation, natural succession commenced on the stabilized sand dunes. The soil in 13-year belt closely resembles to that in sand state, with a predominance of coarse particles and fragile topsoil, whereas the 67-year belt exhibits more advanced soil conditions, nearing a state of ecological stability.

**Sampling design**

We employed the space-for-time substitution method to analyze ecological succession. To evaluate the community composition and spatial development of biological soil crusts (biocrusts), unidimensional interception length measurements conducted along line intercept transects were adopted, hereafter referred to as "transects". For each established sand-binding belt, forty transects, each measuring 1.5 meters in length, were systematically laid out. In consideration of spatial heterogeneity, these transects were positioned at intervals of no less than 2 meters apart. To mitigate potential confounding influences from vegetation, litter, and micro-geomorphological features, the transects were deliberately situated on flat terrain, maintaining a minimum distance of 30 cm from the nearest shrub or tussock and at least 0.5 meters from the nearest soil mound.

Field samplings of biocrust patch were conducted during early (13 years), mid (36 years), and late (67 years) successional stages to compare growth performance of lichen and moss between the centers and edges. Three levels of patch sizes were selected: Small patches: approximately 2 cm in diameter or smaller. Medium patches: consistent with the characteristic scale observed in the late stage, approximately 4 cm in diameter for lichen patches and about 5 cm for moss patches. Large patches: 10 cm or larger in diameter. Sampling was conducted using a 3 cm diameter, 3 cm high PVC ring, collecting on the upper biocrust layer to the laboratory for further analysis. Patch selection was non-random, performed in well-developed, typical areas, ensuring environmental homogeneity by excluding vegetation, mounds, tussocks, and bare patches. Only near-circular patches were chosen, with 9 replicates for each treatment group. These samples were then used for trait-based performance analysis.

**Cellular automaton model**

To operationalize the hypotheses regarding the growth dynamics of biocrust patches, this study employs a spatial model formulated as a probabilistic cellular automaton (CA). The CA model delineates a system where each cell can exist in one of three discrete states, represented by the set,

$$\Sigma=\{S,L,M\}.$$

These states correspond to bare soil ($S$), and soil covered by lichen ($L$) and moss ($M$), respectively. Since the state of cyanobacteria cover is transient and its spatial boundaries are difficult to identify, we combine it with the state of the soil. Furthermore, the model focuses on the development of biocrusts in the inter spaces between plants, thereby excluding the state of vegetation from the simulation. The CA model is structured on a two-dimensional lattice consisting of $I\times I$ cells, with vanishing boundary condition, implying that the lattice’s periphery is perpetually fixed in the soil state. Each cell, indexed by coordinates $r=\{i,j\}$, is of uniform size, denoted by $|r|$. All parameter values for simulation are summarized in the table S1. In our CA model, the behavior of each cell is governed by a set of defined processes:

- **Dispersal:** Vacant cells may sporadically become colonized by biocrusts with a low probability, accounting for air-borne debris or biocrust propagules.
- **Propagation:** Vacant cells adjacent to existing patches possess the potential for colonization. The likelihood of occupancy is positively correlated with the number of neighboring cells, a neighboring facilitation determined by an adjacent range surrounding the focal cell. This correlation indicates the intensity of the short-range positive feedback mechanism.
- **Resource constraint (death):** If a site is occupied by many cells sharing resources, cell death is triggered when local density exceeds the local carrying capacity. Local density is calculated by dividing the number of occupied cells by the total area within the local range.
- **Competition:** In scenarios where both moss and lichen potentially encounter in a cell, mosses are more likely to prevail in spatial competition. This model assumes direct competition rather than indirect exploitation, given that lichen and moss have independent carrying capacities.

We implement these processes in the CA simulation for $t=300$ times, assigning a probabilistic function to each cell: $D(r)$, $P(r)$, $R(r)$, and $C(r)$ (coordinate $r$ is omitted hereafter), corresponding to the occurrence rate of the dispersal, propagation, resource constraint, and competition, respectively. In each iteration, the state of cell is updated according to the transition probability below:

$$\begin{matrix} P\left( S\to L \right) & =P^{L}+D^{L}-\left( P^{L}+D^{L} \right)\cdot\left( P^{M}+D^{M} \right)\cdot C, \\ P\left( S\to M \right) & =P^{M}+D^{M}-\left( P^{M}+D^{M} \right)\cdot\left( P^{L}+D^{L} \right)\cdot\left( 1-C \right) \end{matrix}. [1]$$

They represent the probabilities of lichen and moss colonizing bare soil through dispersal and propagation, respectively, expressed as their individual occupancy rates minus the probability of mutual colonization where the other prevails.

$$\begin{matrix} P\left( L\to S \right) & =d^{L}\cdot R^{L}, \\ P\left( M\to S \right) & =d^{M}\cdot R^{M}, \end{matrix}. [2]$$

Eq. 2 describe the mortality probabilities of lichen and moss patches, triggered by the resource constraint. $d^{L}$and $d^{M}$describe the mortality of lichen and moss with resource constrain.

Furthermore, we also conducted modeling of single-component patch growth to directly verify the hypothesis that asynchronous positive and negative feedbacks leading heavy-tailed to regular PSDs. The single-component model, taking mosses for instance, is simpler as it does not consider competition, and the state transition probabilities are as follows:

$$\begin{matrix} P\left( S\to M \right) & =P^{M}+D^{M} \\ P\left( M\to S \right) & =d^{M}\cdot R^{M} \end{matrix}. [3]$$

In transition equations, $D^{L}$ and $D^{M}$ adopt small and fixed values, representing the success rates of lichen and moss, respectively, randomly colonizing bare soil through dispersal. $P^{L}$ and $P^{M}$ are the probabilities of a cell colonizing other cells through the propagation process. Propagation is achieved through a facilitating mechanism involving adjacent biocrusts. Thus, we define a small circular **adjacent range** with diameters $r_{a}^{L}$ and $r_{a}^{M}$ for lichen and moss, respectively. The strength of the positive feedback is related to the number of adjacent cells of the same type, allowing the propagation intensity to be calculated based on the number of same-type cells within the adjacent range $N_{a}^{L}$ and $N_{a}^{M}$. Consequently, the probability of propagation can be expressed as:

$$P=\left( h+k\cdot N_{a} \right)\cdot\left( 1-R \right), [4]$$

where $h$ and $k$ represent the intercept and slope of the function, respectively, with a high slope indicating strong positive feedback. This function can also employ a higher exponent on $N_{a}$ to achieve more pronounced positive feedback. $R$ represents the state of resource constraint, indicating that the occurrence of propagation presupposes the limited resource constrain.

The resource constraint in the model is described by the number of cells within a large circular range, where biocrust patches compete for resources. This range is referred to as the **local range**, with diameters $r_{l}^{L}$ and $r_{l}^{M}$ for lichen and moss, respectively. The number of lichen and moss cells within this range are denoted as $N_{l}^{L}$ and $N_{l}^{M}$. The local density $\phi^{L}$ and $\phi^{M}$ is calculated as the ratio of these numbers to the total number of cells within the range $N_{l*}^{L}$ and $N_{l*}^{M}$. We define the local carrying capacities for moss and lichen as $K^{L}$ and $K^{M}$, respectively. Therefore, any cell whose local neighbor count exceeds the local carrying capacity results in mortality, represented as:

$$R=\left\{ \begin{matrix} 0 & \text{if }N_{l}>K \\ 1 & \text{if }N_{l}\leq K \end{matrix} \right.. [5]$$

The $K^{L}$ and $K^{M}$ are described by a piecewise unimodal function and a logistic function, since CA model simulated the succession process. These functions capture the increasing trend in carrying capacity of mosses and the initial rise followed by a decline in the carrying capacity of lichens. The trajectories are formulated as follows:

$$\begin{matrix} K^{L} & =\left\{ \begin{matrix} N_{l*}^{L}\cdot\left( \phi_{1}^{L}+\Delta\phi_{1}^{L}\cdot EXP\left( \frac{-\left( t-{100}^{2} \right)}{2\cdot s_{1}^{2}} \right) \right) & If t\leq100 \\ N_{l*}^{L}\cdot\left( \phi_{2}^{L}+\Delta\phi_{2}^{L}\cdot EXP\left( \frac{-\left( t-{100}^{2} \right)}{2\cdot s_{1}^{2}} \right) \right) & If t>100 \end{matrix} \right. \\ K^{M} & =N_{l*}^{M}\cdot\left( \phi_{3}^{M}+\frac{\Delta\phi_{1}^{M}}{1+EXP\left( -\frac{t-100}{s_{2}} \right)} \right) \end{matrix} [6]$$

The parameters $\phi_{1}^{L}$ and $\phi_{2}^{L}$ denote the densities corresponding to initial and final carrying capacities of lichen, respectively. The terms $\Delta\phi_{1}^{L}$ and $\Delta\phi_{2}^{L}$ represent the distances to the peak densities. The parameter $s_{1}$ is the shape parameter for the unimodal function. The parameter $\phi_{3}^{M}$ represents the initial saturated density of moss, and $\Delta\phi_{3}^{M}$ denotes the distance to the plateau density. The parameter $s_{2}$ is the shape parameter for the logistic function. The trajectory parameter values are summarized in the table S2 and the trajectories in simulation are shown in fig. S2. Competition is represented by $C$, which adopts a fixed value, indicating that when moss and lichen encounter each other in the same cell, moss has a higher probability of prevailing.

**Sensitivity analyses and One-component Simulation**

We conducted a sensitivity analysis using a one-component simulation, which compares the evolutionary trajectories of different initial densities (Fig. S3). The results reveal that various initial densities ultimately converge to the same density and spatial pattern. Across all trajectories, two distinct phases are observed: an initial phase of intense fluctuation, followed by a period of stable development into a regular pattern with more defined boundaries. For scenarios starting at low densities, the results of the simulations generally support our working hypothesis that the dominance of positive and negative feedback on patch growth is asynchronous. In the early phase (patch growth phase), characterized by rapid density increases, the positive feedback rule results in a higher likelihood of colonization events occurring in the vicinity of larger patches, thereby promoting the preferential growth of these large patches. This leads to a non-uniform patch size distribution (PSD), which manifests as a heavy-tailed characteristic. As the system transitions into the pattern formation phase, the overall density reaches a plateau and experiences a slight decline. This suggests a spatial optimization of the patches, where larger patches undergo fragmentation due to competitive negative feedback within the dense local environment. This phase is characterized by the attenuation of the heavy-tailed PSD feature, reflecting a transition towards a more regular and evenly distributed patch configuration.

An interesting phenomenon observed in our simulations is the late-stage fragmentation of large patches (as shown in Fig. S3, snapshots 2-3). To investigate this phenomenon, we evaluated the local density ($\phi$) of large patches (those exceeding the average size in a regular pattern), as depicted in Fig. S4. We discovered that when large patches grow in isolation, the center experiences a more intense competition for resources, leading to an inward-to-outward fragmentation (Fig. S4 A). However, when the large patches grow within a regular pattern, their edge faces an additional competition from neighboring patches, resulting in a stronger resource stress (Fig. S4 B). This leads to a contraction rather than fragmentation. Consequently, this explains why the overall biocrust growth of large patches performs poorly in the high density as mentioned in the main text.

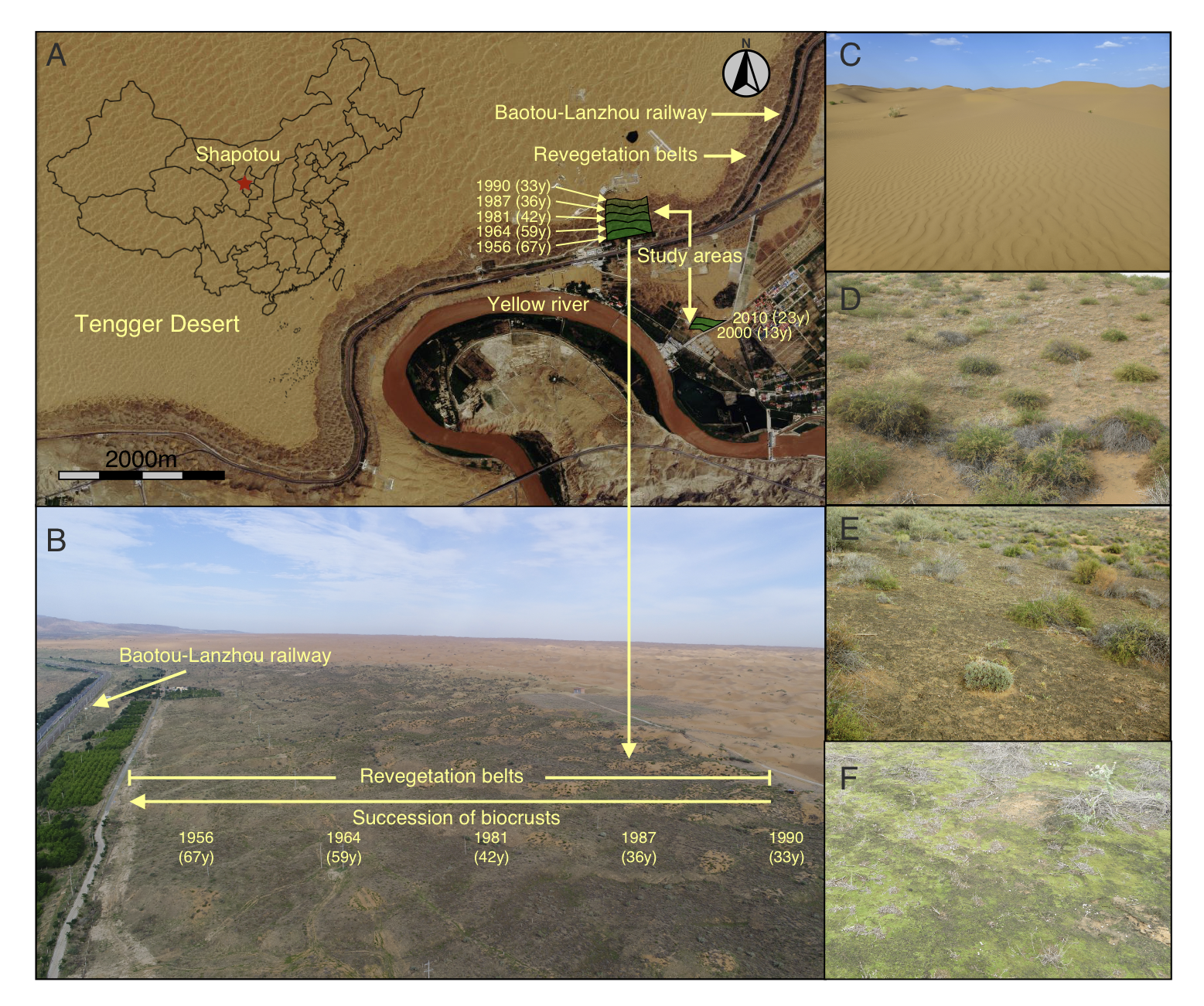

**Fig. S1.** Location and photographs of study areas. A) geographical location of study areas, B) aerial photograph of revegetation chronosequences, illustrating the temporal progression of biocrust development. C) photograph of the bare sand areas that have not yet undergone revegetation, D-F) The change of dominant state of biological crust after vegetation restoration, with cyanobacteria/lichen-nominated biocrusts (D), lichen-nominated biocrusts (E), moss-dominated biocrusts (F).

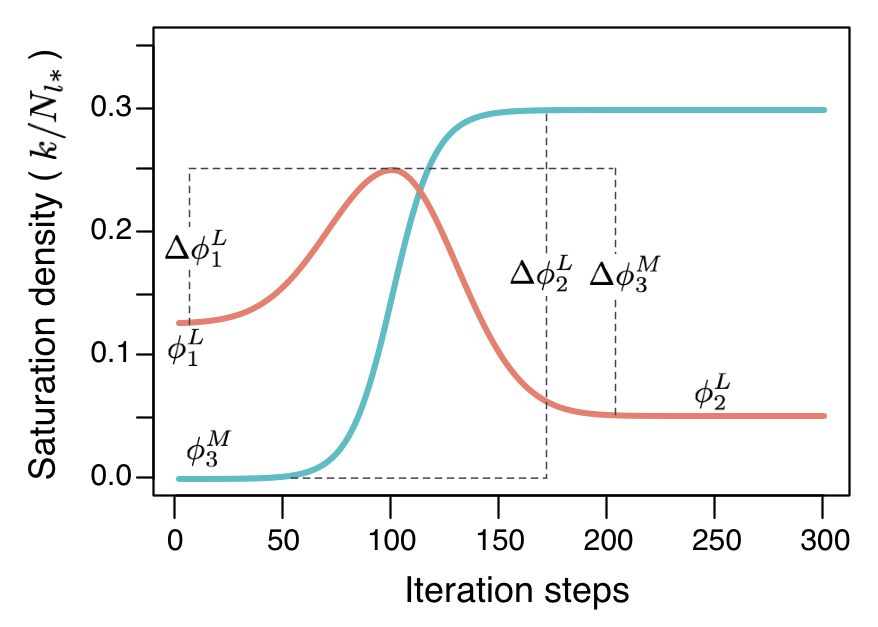

**Fig. S2.** Trajectories of saturated density of lichens (red) and mosses (green), which is denoted as carrying capacity divided by total cell number of local range. These trajectories simulate the dynamic of carrying capacities along succession. The dashed lines are auxiliary to indicate $\Delta\varphi$.

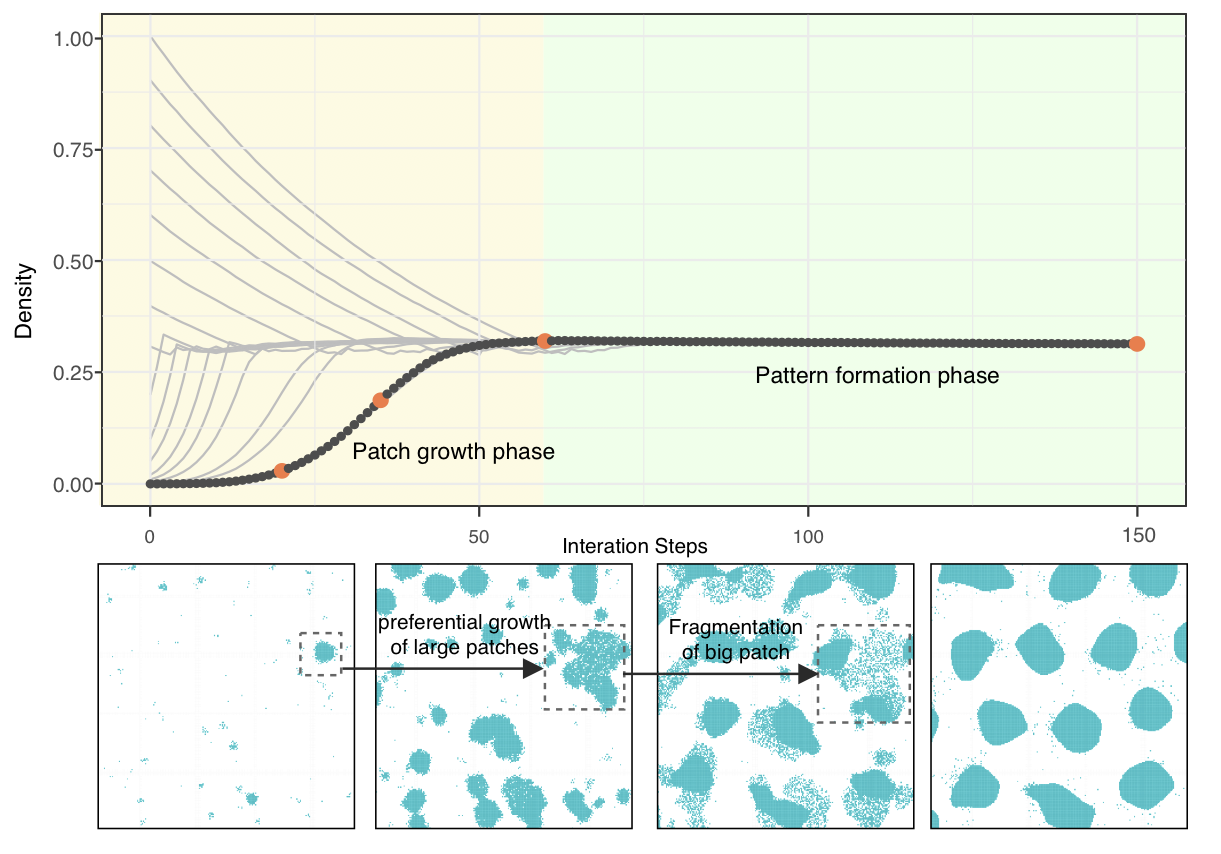

**Fig. S3.** Change of density of simulated patches growth (one component). The snapshots represent the red point. The sensitivity analysis compares the evolutionary trajectories of different initial densities, demonstrating that various initial densities ultimately converge to the same density and spatial pattern. The evolutionary trajectories exhibit two distinct phases: the patch growth phase and the pattern formation phase. During the patch growth phase, density changes rapidly, while in the pattern formation phase, a regular pattern with clear boundaries gradually emerges.

**Table S1.** Parameter list for CA simulation.

| Parameter | Value | Unit | Description |
| --- | --- | --- | --- |
| $I$ | $400$ |  | Gird size of simulation |
| $\vert r\vert$ | $1$ | cm^2^ | Cell size |
| $t$ | $300$ | half year | Iteration times |
| $P^{L}$ & $P^{M}$ | $2.5\times{10}^{-6}$ & $5\times{10}^{-7}$ |  | The probability of lichen and moss occupying soil cell by dispersal |
| $d^{L}$ & $d^{M}$ | $0.95$ |  | The mortality of lichen and moss with resource constrain |
| $r_{a}^{L}$ & $r_{a}^{M}$ | $7$ | cells | Diameters of adjacent range for lichen and moss |
| $h^{L}$ & $h^{M}$ | $0$ & $0.05$ |  | Intercept in relationship of positive feedbacks for lichen and moss |
| $k^{L}$ & $k^{M}$ | $0.01$ & $0.01$ |  | Slope in relationship of positive feedbacks for lichen and moss |
| $r_{l}^{L}$ & $r_{l}^{M}$ | $71$ & $101$ | cells | Diameters of local range for lichen and moss |

**Table S2.** Parameter list for trajectories of local carrying capacity.

| Parameter | Value | Description | Parameter | Value | Description |
| --- | --- | --- | --- | --- | --- |
| $\phi_{1}^{L}$ | 1/8 | Initial saturation density of lichen | $\phi_{3}^{M}$ | 1/10 | Initial saturation density of moss |
| $\phi_{2}^{L}$ | 1/20 | Final saturation density of lichen |  |  |  |
| $\Delta\phi_{1}^{L}$ | 1/8 | Distance to peak density for lichen from initial value | $\Delta\phi_{3}^{M}$ | 1/5 | Distance to plateau density for moss |
| $\Delta\phi_{2}^{L}$ | 1/5 | Distance to peak density for lichen from final value |  |  |  |
| $s_{1}$ | 30 | Shape parameter for unimodal function | $s_{2}$ | 10 | Shape parameter for logistic function |
